# Alternating Magnetic Fields Drive Stimulation of Gene Expression via Generation of Reactive Oxygen Species

**DOI:** 10.1101/2023.08.31.555804

**Authors:** Jordan W. Mundell, Matthew I. Brier, Everest Orloff, Sarah A. Stanley, Jonathan S. Dordick

## Abstract

Magnetogenetics represents a method for remote control of cellular function. Despite successful use of magnetic fields to control gene expression *in vitro* and *in vivo*, the mechanism underlying magnetogenetics is largely unknown, thereby hindering further development and applications. Previous work suggests a chemical-based mechanism involving the generation of reactive oxygen species (ROS) as a platform initiator and downstream signaling molecule. Herein, a chemical biology approach was used to elucidate further the mechanism of radio frequency-alternating magnetic field (RF-AMF) stimulation of a TRPV1-ferritin magnetogenetics platform that leads to Ca^2+^ gating. RF-AMF stimulation of HEK 293T cells expressing TRPV1-Ferritin resulted in a ∼30% and ∼140% increase in intra- and extracellular ROS levels, respectively. Mutations to specific cysteine residues in TRPV1 responsible for ROS sensitivity eliminated RF-AMF driven Ca^2+^-dependent transcription of secreted embryonic alkaline phosphatase (SEAP). Using a non-tethered (to TRPV1) ferritin also eliminated RF-AMF driven SEAP production. These results suggest ferritin-dependent ROS activation of TRPV1 plays a key role in the initiation of magnetogenetics. Furthermore, inhibition of IP_3_R-based endoplasmic reticulum (ER) Ca^2+^ release with Xestospongin C, inhibition of protein kinase C (PKC) activity with Gö 6983, or inhibition of NADPH oxidase (NOX) isoforms 1/2 gp91*^phox^*cytochrome with GSK 2795039 eliminated short-term RF-AMF potentiation of SEAP production with capsaicin. Similarly, Gö 6983 significantly reduced long-term RF-AMF capsaicin potentiation of SEAP production. These results suggest that ROS-activated TRPV1 signaling to increase intracellular Ca^2+^ includes pathways involving PKC, NOX, and the ER. Together, these findings fill in current knowledge gaps in the mechanism of magnetogenetics, which may lead to translational applications in medicine and biotechnology.

## Introduction

External and remote control of signal transduction pathways can provide both on-demand and spatial control over cell function^1–3^ (Arnaud, 2019; Boyden et al., 2005; Gossen et al., 1995). This control can be used to address diseases where aberrant signaling occurs, including cancer, cardiovascular disorders, and neurodegeneration. Several methods to control signaling pathways have been developed, including chemogenetics^4,5^ (Magnus et al., 2011; Gomez et al., 2017) and optogenetics^6,7^ (Gorostiza & Isacoff, 2008; Zhang et al., 2011), that when coupled with recent advances in gene delivery^8^ (Wang et al., 2019) offer potentially new therapeutic routes to address these diseases. Chemogenetics uses designer small molecule drugs specifically to target genetically engineered cell receptors, such as G-coupled protein receptors^9,5^ (Armbruster et al., 2007; Gomez et al., 2017), that regulate cellular pathways when activated, such as neuronal firing and inhibition. However, because chemogenetics requires a pharmacologic intervention, it is strongly dependent on drug pharmacokinetics and pharmacodynamics *in vivo*^10^ (English & Roth, 2015). Optogenetics uses specific wavelengths of light to activate light-sensitive ion channels, such as channelrhodopsin, and provides both spatial and temporal control over signaling pathways that are linked to ion channel activation^2,11,12^ (Boyden et al., 2005; Deisseroth & Hegemann, 2017; Kato et al., 2012). Unfortunately, invasive probes are required *in vivo* due to the poor penetration depth of light^13^ (Nimpf & Keays, 2017).

To avoid these drawbacks, various drug-free and minimally-invasive methods have been developed to control cell signaling, including focused ultrasound^14^ (Pan et al., 2018), electric fields^15,16^ (Lawrence et al., 2022; Krawczyk et al., 2020), and magnetic fields^17–22^ (Huang et al., 2010; Mosabbir & Truong, 2018; Stanley et al., 2012, 2015, 2016; Wheeler et al., 2016). With respect to the latter, Stanley et al.^20^ developed a magnetogenetics platform consisting of an engineered transient receptor potential vanilloid 1 channel (TRPV1 or V1) fused with an anti-green fluorescent protein (GFP)-nanobody (αGFPnb) that interacts with engineered ferritin that contains an integrated GFP-tagged ferritin dimer (FtD) subunit, herein referred to as αGFPnb-V1/GFP-FtD. TRPV1 is a well characterized cation channel that selectively gates Ca^2+^ ions via a number of activation mechanisms, including temperature^23,24^ (Caterina et al., 1997; Grandl et al., 2010), mechanical force^25^ (Gaudet, 2008), reactive oxygen species (ROS)^26^ (Ogawa et al., 2016), voltage^24^ (Grandl et al., 2010), and selective ligands such as capsaicin^23^ (Caterina et al., 1997). In the presence of a radio frequency alternating magnetic field (RF-AMF), the TRPV1-ferritin complex (αGFPnb-V1/GFP-FtD) is activated, leading to Ca^2+^ influx^27^ (Brier et al., 2020). Increased intracellular calcium is then used to control Ca^2+^-dependent gene transcription via a synthetic Ca^2+^-dependent promotor system consisting of triplicate serum response elements (SRE)^28^ (Miranti et al., 1995), adenosine 3′,5′-cyclic monophosphate (cAMP) response elements (CRE)^29^ (Grewal et al., 2000), and nuclear factor of activated T-cells response elements (NFATRE)^30^ (M.G. Pan et al., 2013) all upstream of a minimal promoter (minP)^27^ (Brier et al., 2020). Specifically, RF-AMF was used to express secreted embryonic alkaline phosphatase (SEAP) *in vitro*^19,20,27^ (Stanley et al., 2012, 2015; Brier et al., 2020) as well as proinsulin to reduce blood glucose levels *in vivo*^20^ (Stanley et al., 2015). Tethering of the engineered ferritin to TRPV1 is critical, as TRPV1 activation in the RF-AMF is minimal for non-tethered cytosolic ferritin (absence of the anti-GFP nanobody) or myristoylated ferritin that inserted ferritin randomly into the cell membrane^20^ (Stanley et al., 2015).

Despite significant interest in exploiting magnetic fields to control gene expression, the precise mechanism of RF-AMF driven magnetogenetics remains largely unknown. Early hypotheses involving thermal or mechanical activation of TRPV1 in the presence of RF-AMF were considered improbable due to ferritin’s iron oxide core being unlikely to generate sufficient heat or mechanical motion at a sufficient distance from the protein core to influence TRPV1^31^ (Meister, 2016). However, depending on the physicochemical properties of ferritin’s iron oxide core, sufficient localized heating is theoretically possible^32^ (Barbic, 2019). Recently, Hernández-Morales et al.^33^ (2020) proposed a chemical mechanism for RF-AMF stimulation of a ferritin-conjugated TRPV1/4 magnetogenetics platform via lipid oxidization due to generation of localized ROS. Brier et al.^27^ (2020) also proposed a chemical mechanism of RF-AMF driven activation of αGFPnb-V1/GFP-FtD, and critically, demonstrated the importance of ROS as a signaling molecule in magnetic field-driven gene expression. Activation of this cellular construct was dependent on magnetic field frequency and field strength, and magnetic activation was potentiated in the presence of the TRPV1 agonist capsaicin, even at temperatures well below capsaicin-driven channel gating in the absence of the magnetic field^27^ (Brier et al., 2020). These studies confirmed that TRPV1 activation was necessary for RF-AMF Ca^2+^ channel gating as addition of the TRPV1 antagonist, AMG-21629, prevented TRPV1 activation by RF-AMF, as reflected in loss of SEAP production, with and without capsaicin. Addition of N-acetylcysteine also eliminated SEAP production confirming a key role for ROS in RF-AMF TRPV1 activation. Capsaicin potentiation was reduced in the presence of apocynin, a broad NADPH oxidase (NOX) isoforms 1 and 2 inhibitor; however, TRPV1 activation was not affected, suggesting that capsaicin potentiation likely resulted from inflammatory signaling leading to NOX1/2 activation and Ca^2+^ ion release. Brier et al.^27^ (2020) concluded by hypothesizing a secondary pathway being activated due to increased Ca^2+^ levels from RF-AMF/capsaicin potentiation. This pathway involves Ca^2+^-dependent activation of protein kinase C (PKC) leading to NOX assembly, resulting in increased ROS levels and Ca^2+^ release from the endoplasmic reticulum (ER).

These results provide tantalizing insight into the mechanism of magnetogenetics, yet the precise activation of TRPV1 in the presence of RF-AMF remains elusive. Elucidating this mechanism in detail may allow further development of magnetogenetic tools to modulate cell signaling and could drive new applications of remote cellular control. In the current work, we performed a series of biomolecular studies, including genetic modification of the TRPV1, to elucidate further the mechanism of TRPV1 activation in an RF-AMF and the underlying mechanisms of the potent potentiation of Ca^2+^ ion signaling in the presence of capsaicin and for the downstream gene activation leading to specific reporter protein expression. As a result, we identify the precise molecular role of ROS in the mechanism of magnetogenetics.

## Results

### Chemical inhibition of magnetogenetics pathways

To study the Ca^2+^ signaling cascade directly and indirectly downstream of RF-AMF activation of TRPV1, we used the same Ca^2+^-dependent SEAP (Ca-SEAP) construct as in our prior work^27^ (Brier et al., 2020) (**Figure 1A**). HEK 293T cells stably expressing the αGFPnb-V1/GFP-FtD magnetogenetics platform and transfected with Ca-SEAP treated with RF-AMF (∼501 kHz, 27 mT) displayed significant SEAP production following 2 h stimulation, 2.3 ± 0.3 -fold over baseline levels (**Figure 1B**). Thus, 2 h stimulation blocks were used throughout this study except where indicated. Four pathway inhibitors involved in key steps in Ca^2+^-dependent signaling, particularly involving ROS generation, were used to parse the contributing factors in RF-AMF driven activation of TRPV1: (i) xestospongin C (XC), an inhibitor of sarcoplasmic/endoplasmic reticulum ATPase that blocks 1,4,5-trisphosphate (IP_3_)-driven Ca^2+^ release from the ER^34^ (Oka et al., 2002); (ii) ryanodine (Rya), a ryanodine receptor (RyR) antagonist that blocks RyR-based Ca^2+^ release from the ER-based intracellular Ca^2+^ stores^35^ (Pisaniello et al., 2003); (iii) Gö 6983 (Gö, 3-[1-[3-(dimethylamino)propyl]-5-methoxy-1H-indol-3-yl]-4-(1H-indol-3-yl)-1H-pyrrole-2,5-dione), a broad-spectrum PKC inhibitor^36^ (Young et al., 2005); and (iv) GSK 2795039 (GSK, N-(1-isopropyl-3-(1-methylindolin-6-yl)-1H-pyrrolo[2,3-b]pyridin-4-yl)-1-methyl-1H-pyrazole-3-sulfonamide), a generally selective inhibitor of the NOX gp91*^phox^* cytochrome^37^ (Hirano et al., 2015).

**Figure 1:**
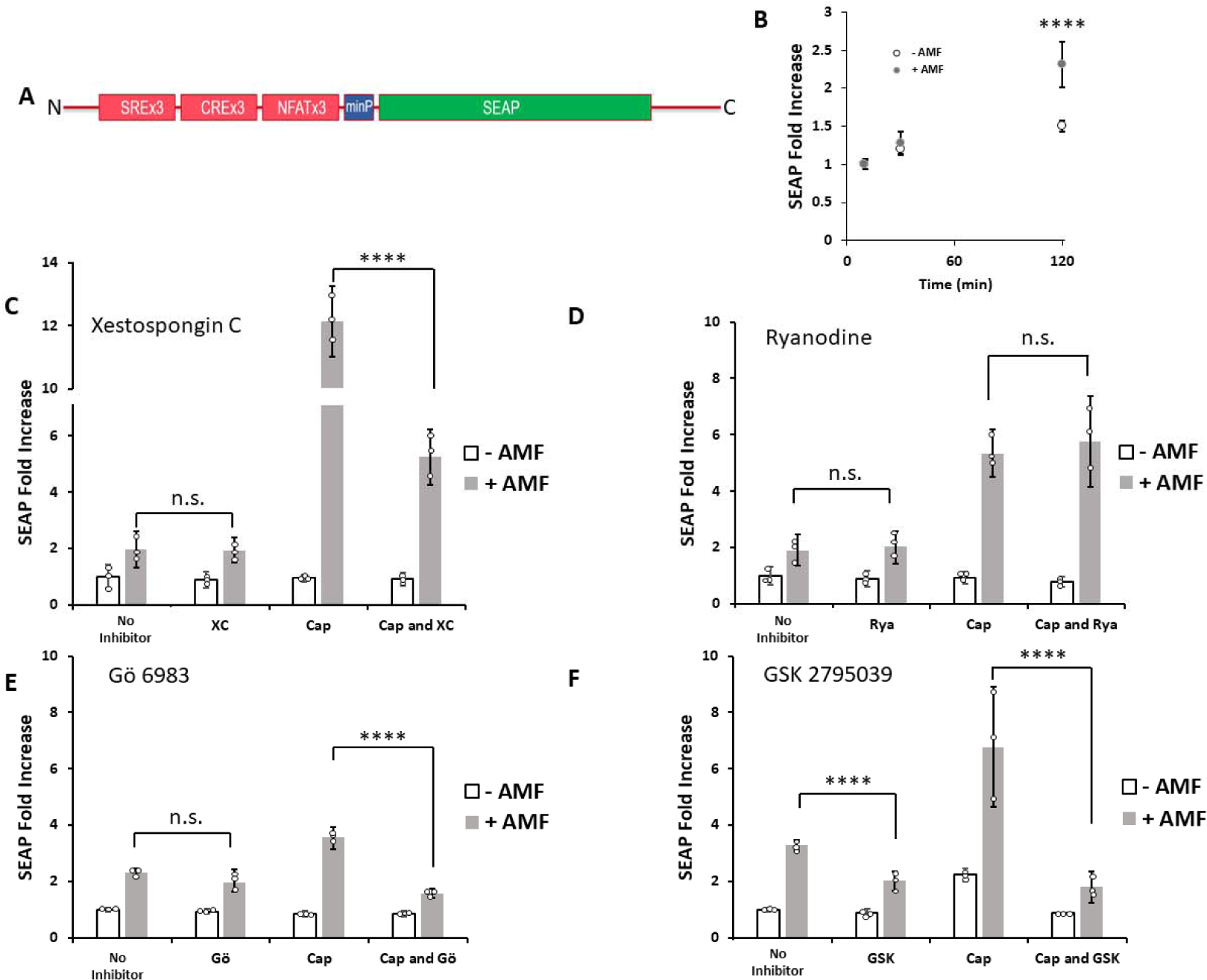
Inhibition of magnetic field-driven TRPV1 activation. HEK-293T cells stably expressing the αGFPnb-V1/GFP-FtD construct and transfected with Ca^2+^-dependent SEAP gene were stimulated with RF-AMF (501 kHz, 27.1 mT) for 2 h with ± 1.0 mM capsaicin (Cap) and ± indicated inhibitors. (A) Ca^2+^-dependent SEAP construct comprised of triplicate of SRE, CRE, and NRE in conjunction with the minimal promoter. (B) SEAP production time course. (C) Xestospongin C, XC, 1.0 μM. (D) Ryanodine, Rya, 1.0 μM. (E) Gö 6983, Gö, 5.0 μM. (F) GSK 2795039, GSK, 100 μM. (C-F) All +AMF groups were statistically significant over respective-AMF groups (p < 0.001, not shown). N = 3 for all treatments, mean ± S.D. **** (p < 0.0001).

HEK 293T cells stably expressing the αGFPnb-V1/GFP-FtD magnetogenetics construct and transiently transfected with a Ca-SEAP gene were exposed to RF-AMF (∼501 kHz, 27 mT) for 2 h in the presence or absence of 1.0 μM capsaicin and the presence or absence of an indicated inhibitor. XC at 1.0 μM exerted no significant effect on RF-AMF driven SEAP production (1.9 ± 0.7 and 2.0 ± 0.5 fold over baseline without and with XC, respectively); however, a significant decrease was observed for RF-AMF/Cap potentiation (12.1 ± 1.1 and 5.3 ± 1.0 fold without and with XC, respectively; p < 0.0001) (**Figure 1C**). The addition of 1.0 μM Rya had no significant effect on SEAP production with either RF-AMF or RF-AMF/Cap potentiation (**Figure 1D**). Together these results suggest that ER-based Ca^2+^ release is only significant in the RF-AMF/Cap potentiation pathway, and not in RF-AMF alone, due to IP_3_ receptor (IP_3_R) activation and not RyR response.

To examine the contribution of PKC signaling in RF-AMF TRPV1 activation and its potentiation by capsaicin, we examined the effects of a broad-spectrum PKC inhibitor, Gö. No significant effect on RF-AMF driven SEAP production in the absence of Cap was observed in the presence of 5.0 μM Gö (2.3 ± 0.2 and 2.0 ± 0.3 fold over baseline without and with Gö, respectively), while RF-AMF/Cap potentiation was essentially abolished (3.6 ± 0.4 vs. 1.6 ± 0.2 fold over baseline without and with Gö, respectively, p < 0.0001), with SEAP levels similar to that in the absence of Cap (**Figure 1E**). This suggests that PKC inhibition prevents capsaicin potentiation of downstream calcium signaling, likely including NOX assembly^37^ (Hirano et al., 2014) and ER Ca^2+^ release^38^ (Sakurada et al., 2019). Finally, addition of 100 μM GSK, a selective inhibitor of NOX gp91*^phox^*cytochrome, resulted in a significant decrease in SEAP production under both RF-AMF and RF-AMF/Cap conditions (**Figure 1F**). Specifically, RF-AMF driven SEAP production dropped from 3.3 ± 0.2 to 2.0 ± 0.3 fold over baseline without and with GSK, respectively (p < 0.0001), while RF-AMF/Cap potentiation dropped from 6.8 ± 2.1 to 1.8 ± 0.6 fold over baseline without and with GSK, respectively (p < 0.0001). The inhibition of SEAP production by GSK in either the RF-AMF or RF-AMF/Cap conditions indicates that NOX1/2 plays a significant role in RF-AMF driven TRPV1 activation.

### Longer-term and multiple AMF stimulation

To understand the effect of longer-term AMF stimulation on SEAP production, HEK-293T cells stably expressing the αGFPnb-V1/GFP-FtD construct and transfected with the Ca-SEAP gene were stimulated with (+) and without (−) RF-AMF (501 kHz, 27.1 mT) for a sequence of three 2 h treatment blocks. This resulted in five treatment sequences: (− − −), (+ + +), (+ + −), (+ − +), and (+ − −), indicating whether RF-AMF (+) or no RF-AMF (−) was used for each 2 h block with SEAP values presented as fold relative to initial 2 h baseline (- AMF) production (**Figure 2**). Baseline SEAP production, i.e. (− − −), remained relatively constant for each 2 h treatment. Repeated AMF stimulation (+ + +) resulted in accelerated production, with each subsequent treatment producing significantly more SEAP than the previous treatment (1.6 ± 0.2, 2.2 ± 0.3, and 3.4 ± 0.4 fold, respectively, p < 0.0001) (**Figure 2A**). This suggests that magnetogenetics can be sustained for regulation of gene expression over many hours.

**Figure 2:**
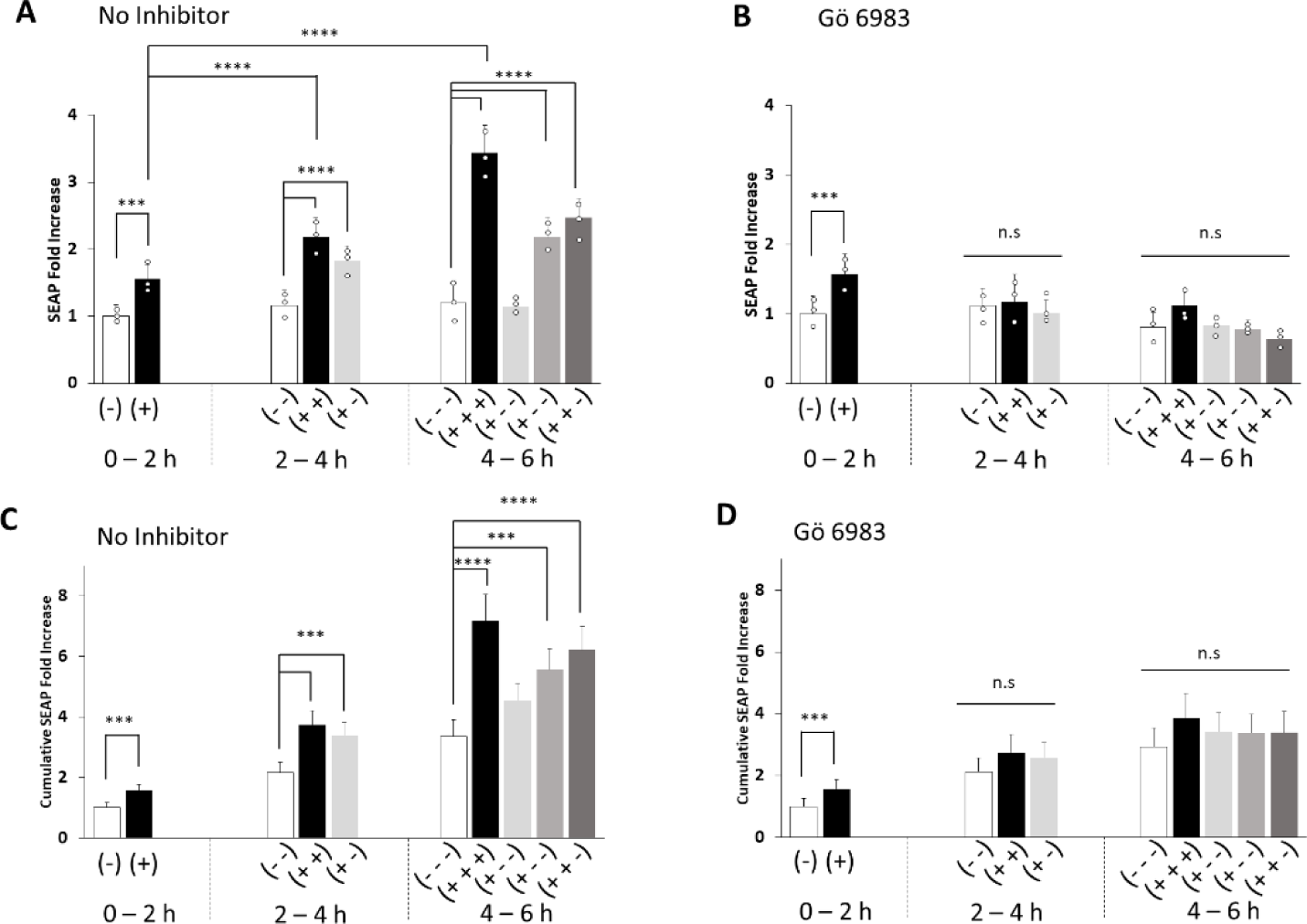
Long-term and multiple RF-AMF stimulation. HEK-293T cells, stably expressing the aGFPnb-V1/GFP-FtD construct and transfected with Ca2+-dependent SEAP gene, were stimulated with ± RF-AMF (501 kHz, 27.1 mT) in 2 h treatment blocks for up to 6 h. Dashed and solid lines indicate (−) and (+) AMF for a 2 h block, respectively. Time-point SEAP production for (A) no inhibitor and (B) + 5.0 mM Gö 6983. Cumulative SEAP production for (C) no inhibitor and (D) + 5.0 mM Gö 6983. Treatment conditions are: no AMF (− − −), AMF for 6 h (+ + +), AMF for 4 h followed by no AMF for 2 h (+ + −), AMF for 2 h followed by no AMF for 2 h followed by AMF for 2 h (+ − +), and AMF for 2 h followed by no AMF for 4 h (+ − −). For all cases, N = 3, mean ± S.D.

Next, we examined whether the downstream calcium signaling pathways recruited by prolonged RF-AMF stimulation of TRPV1 differ from those with short term stimulation. Specifically, we examined the effects of Gö on longer term RF-AMF SEAP production, as PKC is critical in activation of NOX due to phosphorylation of p47*^phox^* ^39^(Cosentino-Gomes et al., 2012), and consequently a key initiator of the entire PKC/NOX/ER pathway. The inclusion of 5.0 μM Gö did not affect SEAP production after 2 h of RF-AMF stimulation (1.6 ± 0.3 fold); however, it did significantly lower longer-term SEAP production due to sequential RF-AMF stimulations as shown by the drop in production from 3.4 ± 0.4 (**Figure 2A**) to 1.1 ± 0.4 fold after the third RF-AMF treatment (+ + +) (**Figure 2B**) normalized to the 2 h no AMF time point (p < 0.0001). These trends are exacerbated with respect to cumulative SEAP production. In the absence of Gö, total SEAP production for (+ + +) increased to 7.2 ± 0.9 fold after 6 h compared to the baseline value of 3.4 ± 0.4, p < 0.0001 (**Figure 2C**). With Gö, (+ + +) drops significantly to 3.5 ± 0.3, p < 0.0001 (**Figure 2D**). The increased SEAP production and inhibition via Gö mirrors the trends seen from RF-AMF/Cap potentiation, suggesting that prolonged RF-AMF stimulation may activate the PKC/NOX/ER pathway, however, without the need for capsaicin potentiation. Due to the relatively weak response from RF-AMF compared to capsaicin, longer treatment times likely are needed to build up sufficient cytosolic Ca^2+^ levels to activate this pathway.

Interestingly, for the (+ − −) treatment, after the initial 2 h of RF-AMF stimulation, SEAP production still occurs at elevated levels (1.8 ± 0.2 fold) similar to the original RF-AMF stimulation (1.6 ± 0.2 fold) during the subsequent 2 h no-treatment block. SEAP production during the final 2 h no-treatment, however, dropped back to baseline levels (1.1 ± 0.1 fold) (**Figure 2A**). This suggests that RF-AMF stimulation has a lingering impact on SEAP production, but will return to baseline levels after about 2 h.

These findings are further corroborated when AMF stimulation was altered from a continuous stimulation to pulsed stimulations (**Figure S1**). When AMF was pulsed with equal on and off phases over 4 h (i.e., 2 h of +AMF and 2 h of – AMF total), a positive correlation was obtained between pulse duration and both SEAP (**Figure S1A**) and ROS (**Figure S1B**) production. This indicates that despite total AMF stimulation being constant, a mechanistic difference was observed, in which prolonged AMF stimulation can produce sufficient ROS levels to continue to activate the platform, even after AMF is removed.

### Quantification of reactive oxygen species

The significant inhibition of RF-AMF driven SEAP production by GSK, with and without capsaicin, suggests that NOX1/2-generated ROS in HEK-293T cells, plays a role in TRPV1 activation, and this ROS is potentiated in the presence of capsaicin. However, even in the absence of capsaicin, ROS may be formed from ferritin upon RF-AMF treatment without a potentiator^33^ (Hernandez-Morales et al., 2020), likely due to AMF effecting local iron levels around ferritin^40,41^ (Céspedes et al., 2009, 2010). To understand better the role of ROS under RF-AMF conditions, HEK-293T cells stably expressing the αGFPnb-V1/GFP-FtD magnetogenetics construct were exposed to RF-AMF and both intra- and extracellular ROS levels were measured using DCFDA and Amplex Red, respectively. RF-AMF stimulation for 2 h resulted in a 1.3 ± 0.2 fold increase in intracellular ROS production over baseline, p < 0.001 (**Figure 3A**). *Tert*-butyl hydrogen peroxide (TBHP, 50 μM) was used as a positive control. Similarly, 2 h RF-AMF stimulation resulted in a 2.4 ± 1.0 fold in extracellular ROS over baseline, p < 0.001 (**Figure 3B**). Capsaicin alone resulted in 2.5 ± 0.6 fold over baseline, p < 0.0001, while the RF-AMF/Cap potentiation results mirrored the SEAP production, increasing to 5.3 ± 1.3 fold over baseline **(Figure 3B**). Furthermore, RF-AMF driven extracellular ROS production increased throughout the 2 h incubation (**Figure 3C**), becoming statistically significant at 30 min (p < 0.0001). Ferritin-derived ROS will be entirely intracellular while NOX-derived ROS has been shown to increase both intra- and extracellular ROS. However, increased ROS levels within the cell would also be able to diffuse through the cell membrane into extracellular space, thus, intra- and extracellular ROS generation is interconnected. These results indicate that RF-AMF is capable of generating both intra- and extracellular ROS that is consistent with activation of TRPV1 and Ca^2+^ channel gating.

**Figure 3:**
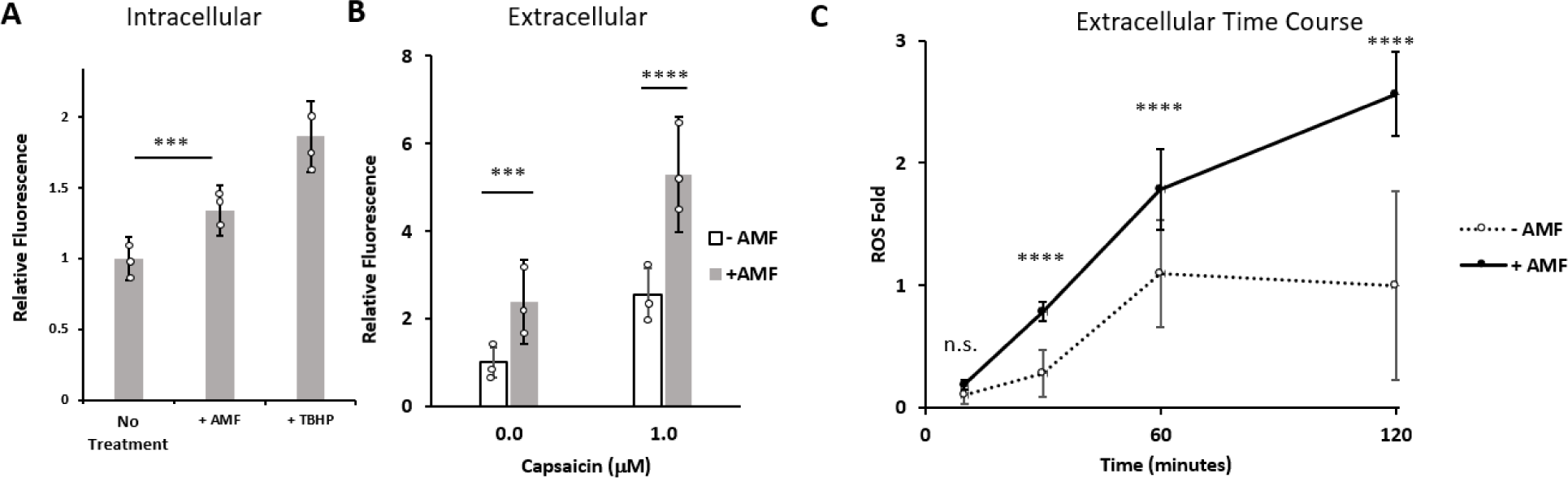
RF-AMF induced ROS detection. HEK-293T cells, stably expressing the αGFPnb-V1/GFP-FtD construct, were stimulated with RF-AMF (501 kHz, 27.1 mT) and intra-an extracellular ROS production were determined using Amplex Red/horseradish peroxidase and DCFDA, respectively. (A) Intracellular ROS formation after 2 h of RF-AMF stimulation. TBHP (50 mM, *tert*-butyl hydrogen peroxide was used as positive control). N = 6. (B) Extracellular ROS formation after 2 h of RF-AMF stimulation with and without 1.0 mM capsaicin present. N = 3. (C) Time course for extracellular ROS formation over 2 h of RF-AMF stimulation. Data reported as fold relative to baseline 2 h incubations. N = 3. Mean ± S.D.

The effect of the previously mentioned inhibitors (XC, Rya, Gö, and GSK) on ROS generation with and without AMF stimulation was also assessed (**Figure S2**). Both XC and Rya had no effect on ROS generation, either baseline or from AMF stimulation. GSK resulted in a significant decrease in AMF-induced ROS generation, causing a drop from 1.7 ± 0.5 fold to 1.1 ± 0.2 fold over baseline (p < 0.05). Similarly, Gö reduced ROS levels to 1.3 ± 0.4 fold (however, non-significantly). GSK and Gö also significantly reduced ROS levels in the absence of AMF (dropping to 0.6 ± 0.3 and 0.6 ± 0.2 fold, respectively, p < 0.05) (**Figure S2**). These results suggest that GSK and Gö have significant effects on ROS levels, both in the presence and absence of AMF, likely due to interactions with NOX, while XC and Rya do not.

### TRPV1 engineering

The activation of TRPV1 due to ROS has been hypothesized to be due to the oxidation of specific cysteine residues. Specifically, Ogawa et al.^26^ (2016) identified two cysteine residues in TRPV1 that become oxidized in the presence of ROS. To assess whether RF-AMF driven ROS production impacts TRPV1 channel gating, mutated magnetogenetics constructs were created, replacing cysteine residues with serine at the two sites indicated in Ogawa et al.^26^ (2016). As shown in **Figure 4A**, one site (C257) is located on one of the N-terminal ankyrin domain loops, while the other is near the C-terminus (C741). Three TRPV1 mutants within the αGFPnb-V1/GFP-FtD construct were prepared and examined in the presence and absence of RF-AMF – the C257S and C741S single mutants and the C257S/C741S double mutant.

**Figure 4:**
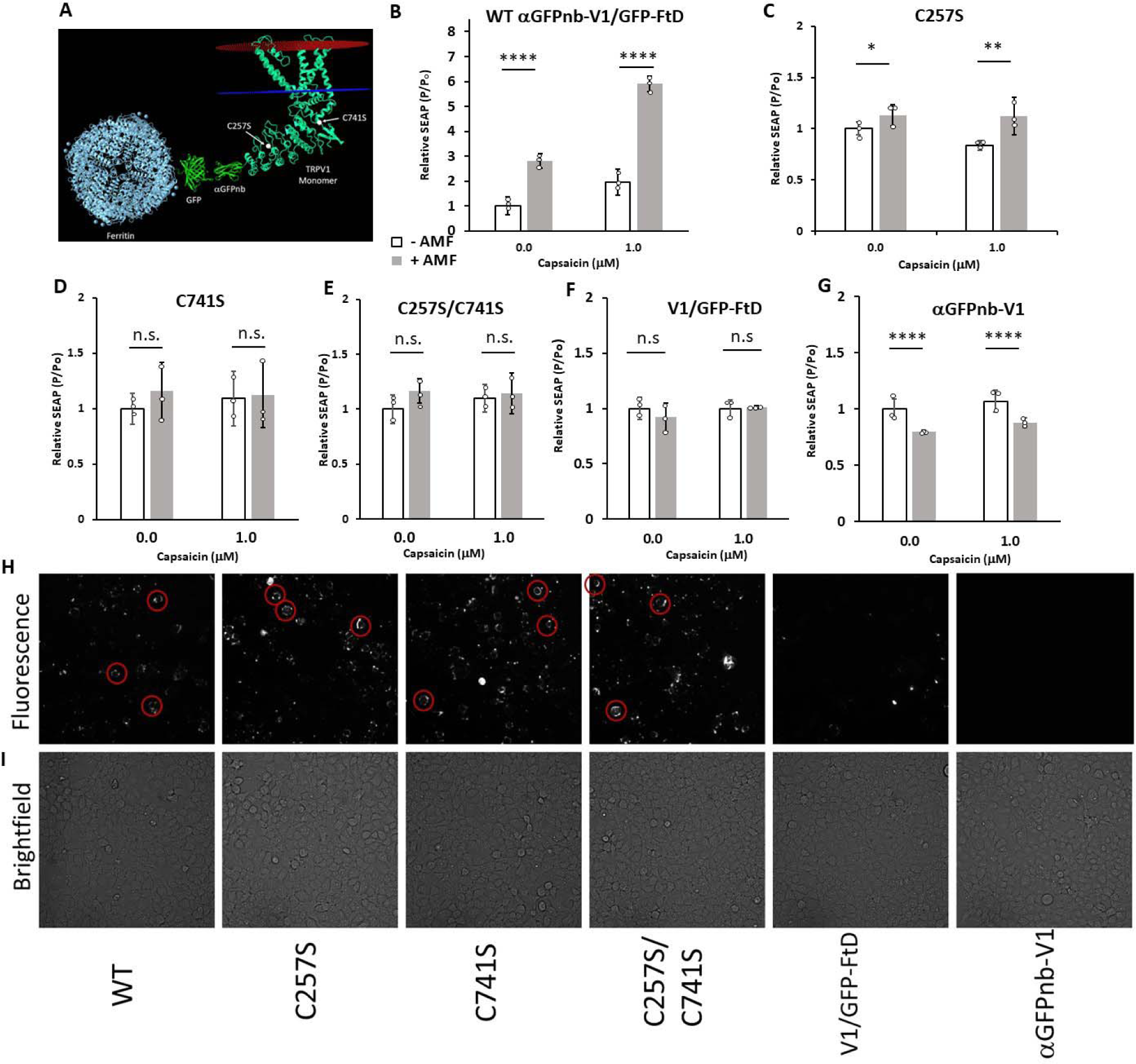
Engineering of the magnetogenetics platform. Several genetic modifications were made to the αGFPnb-V1/GFP-FtD magnetogenetics construct. (A) Schematic of the ferritin-tethered TRPV1 indicating the locations of C257 and C741 residues. (B) Wild-type TRPV1 within the αGFPnb-V1/GFP-FtD construct, transfected with Ca^2+^-dependent SEAP construct and stimulated with RF-AMF (501 kHz, 27.1 mT) for 2h. For (C-G), all platforms were independently transfected with the αGFPnb-V1/GFP-FtD construct along with Ca^2+^-dependent SEAP into HEK-293T cells and subsequently stimulated with RF-AMF (501 kHz, 27.1 mT) for 2 h. (C) C257S mutation, (D) C741S mutation, (E) C257S/C741S double mutation, (F) V1/GFP-FtD (removal of anti-GFP nanobody), and (G) aGFPnb-V1 (removal of GFP-tagged ferritin dimer). N = 3 for all treatments, mean ± S.D. (H-I) HEK 293T cells were transfected with aGFPnb-V1/GFP-FtD (WT) or indicated engineered platforms. (H) GFP fluorescence and (I) Brightfield images were acquired on a Thermo Scientific Cellomics ArrayScan XTI at 20x magnification. Red circles depict ring-like fluorescence, indicating expression of the platform and recruitment to the cell membrane.

HEK-293T cells were transiently transfected with WT and mutant αGFPnb-V1/GFP-FtD TRPV1 channels, and SEAP production determined with and without exposure to RF-AMF for 2 h. Wild-type αGFPnb-V1/GFP-FtD showed expected increases in SEAP production, with RF-AMF and RF-AMF/Cap potentiation increasing to 2.8 ± 0.3 fold and 5.9 ± 0.3 fold over the baseline condition, respectively, p < 0.0001 (**Figure 4B**). The C257S mutant caused muted sensitivity to RF-AMF with or without capsaicin (**Figure 4C**). The C741S mutant was insensitive to RF-AMF driven SEAP production with and without capsaicin (**Figure 4D**), as was the C257S/C741S double mutant (**Figure 4E**). Significantly higher ROS levels were still observed, p< 0.05 (**Figure S3**), confirming these Cys residues are crucial for platform stimulation. These results, together with Ogawa et al.^26^ (2016) support the hypothesis that ROS generated by ferritin in the presence of a RF-AMF activates TRPV1 leading to channel gating and this activation is unable to occur when the two susceptible Cys residues are eliminated.

The αGFPnb-V1/GFP-FtD construct was also modified to eliminate the ability of ferritin to be tethered to TRPV1. To this end, removing GFP-FtD or the anti-GFP nanobody bound to the N-terminus of TRPV1 (αGFPnb) from the construct would be expected to prevent ferritin from binding to the ion channel. HEK-293T cells were transiently transfected with either the construct without the anti-GFP nanobody (V1/GFP-FtD, **Figure 4F**) or without the GFP-FtD (αGFPnb-V1, **Figure 4G**) and the Ca-SEAP construct, and then they were treated with and without RF-AMF (501 kHz, 27 mT) and Cap for 2 h. The absence of the αGFPnb abrogated RF-AMF induced SEAP production (0.9 ± 0.1 fold vs. 1.0 ± 0.1 fold), regardless of capsaicin treatment (1.0 ± 0.1 vs. 1.0 ± 0.0 fold relative to baseline, respectively) (**Figure 4F**). Similar results were obtained when the GFP-FtD was removed, with RF-AMF induced production eliminated (**Figure 4G**). Similar to the ROS-insensitive mutants, ROS levels were still significantly higher for the V1/GFP-FtD and αGFPnb-V1 platforms when exposed to AMF (with αGFPnb-V1 having a relatively smaller increase due to the removal of exogenous ferritin) (**Figure S3**). These results confirm the critical role of ferritin tethering to TRPV1 in the proximal generation of ROS to the channel.

To confirm expression and functionality of engineered TRPV1, fluorescent and brightfield images (**Figure 4H** and **Figure 4I**) and responsiveness to capsaicin (**Figure S4**) were assessed. All three ROS-insensitive mutants exhibited similar fluorescence patterns to the WT, with rings of GFP fluorescence visible (emphasized with red circles in the figure), indicating proper assembly of the platform in the cell membrane (**Figure 4H**). Removal of the αGFPnb prevents ferritin conjugation with TRPV1. However, since GFP-tagged ferritin is still present, diffuse GFP is still detected, but with lack of the aforementioned rings (**Figure 4H**). Similarly, all modified platforms experienced significant SEAP production in response to a capsaicin gradient (p < 0.05), indicating TRPV1 functionality in all platform variants (**Figure S4**). Note that differences in SEAP response to capsaicin in **Figure S4** from that in **Figure 4** is due to the different temperatures and durations (37°C for 3 h for **Figure S4** vs. 32°C for 2 h for **Figure 4**).

## Discussion

The field of magnetogenetics has been controversial and has led to significant skepticism in the literature^31^ (Meister, 2016), despite the fact that the effectiveness of magnetogenetics has been demonstrated both *in vitro* and *in vivo* in multiple publications and by multiple independent research groups^19–22,27,33^ (Brier et al., 2020; Hernandez-Morales et al., 2020; Stanley et al., 2012, 2015, 2016; Wheeler et al., 2016). This skepticism cannot be resolved without the elucidation of a precise mechanism of magnetic field activation of cell signaling, for example, ferritin-mediated activation of TRPV1 Ca^2+^ influx and resulting Ca^2+^ signaling. Initial hypotheses on the mechanism of magnetic field-driven activation of TRPV1 included thermal and mechanical contributions, based on known phenomena of magnetic nanoparticles in AMFs^42^ (Lima et al., 2013), which would provide the necessary internal energy required to gate the channel. However, theoretical calculations based on the physical properties of ferritin have called those hypotheses into question^31^ (Meister, 2016). Nonetheless, RF-AMF is known to affect iron levels within ferritin^40,41^ (Céspedes et al., 2009; Céspedes & Ueno, 2010), which could be a driving force behind localized ROS production^33^ (Hernandez-Morales et al., 2020), and this has been demonstrated to be strongly dependent on field frequency and strength^27^ (Brier et al., 2020). Lack of a functional mechanism impacts ultimate biomedical applications and further optimization of magnetogenetic tools. For this reason, we embarked on a more detailed elucidation of the TRPV1-ferritin magnetogenetics mechanism.

Our study is consistent with ROS-driven TRPV1 channel gating. When exposed to RF-AMF, the ferritin in the tethered αGFPnb-V1/GFP-FtD TRPV1 construct will locally produce ROS (**Figure 3**) in the vicinity of TRPV1. We propose that the presence of a sufficient amount of ROS, such as formation of H_2_O_2_, will result in oxidation of susceptible cysteine residues in TRPV1, consistent with physiological activation of TRPV1 via ROS^43^ (Taylor-Clark, 2016). This increased ion channel sensitivity is consistent with the work of Ogawa et al.^26^ (2016) who demonstrated that C257 and C741 on TRPV1 are oxidized in the presence of H_2_O_2_, which enables TRPV1 to gate at lower temperatures than the normal 42°C. This is also consistent with other orthogonal stimuli regulating TRPV1 activation, e.g., RF-AMF reducing the thermal threshold for capsaicin-driven TRPV1 gating^24^ (Grandl et al., 2010). In the absence of tethered ferritin, the local concentration of ROS in the TRPV1 pore vicinity may be too low to activate the ion channel. This result is consistent with Stanley et al.^20^ (2015) who observed similar effects with myristoylated ferritin versus directly bound nanoparticles. Interestingly, the C741S mutation appears to reduce RF-AMF TRPV1 activation more significantly than the C257S mutation (**Figures 4C and D**), perhaps due to the former being closer to the pore domain of the TRPV1 channel.

We previously hypothesized that activation of TRPV1 by RF-AMF appeared distinct from the mechanism of capsaicin potentiation due to inhibitors (apocynin, AMG-21629, and N-acetylcysteine) having no effect on AMF alone, but significantly reducing or eliminating RF-AMF/Cap potentiation^27^ (Brier et al., 2020). We suggested that RF-AMF generated ROS in conjunction with capsaicin would reduce the thermal threshold of TRPV1, hypersensitizing the channel, and resulting in increased Ca^2+^ influx compared to RF-AMF alone. The heightened intracellular Ca^2+^ levels could then activate various Ca^2+^/ROS pathways such as PKC-activation of NOX1/2, resulting in addition ROS generation and subsequently ER-Ca^2+^ release, hence a PKC-NOX-ER feedback loop. While it is likely this RF-AMF/Cap potentiation pathway still occurs when capsaicin is present, our results herein (**Figure 2**) suggest that AMF alone is capable of stimulating the PKC-NOX-ER feedback loop. However, since RF-AMF stimulation is weaker than capsaicin, it requires longer durations of RF-AMF stimulation to build up sufficient intracellular Ca^2+^ to initiate this secondary pathway. This hypothesis is further supported by SEAP and ROS production positively correlating with RF-AMF pulse duration, indicating that increased exposure to RF-AMF can produce sufficient levels of ROS to continue platform stimulation even after RF-AMF is removed (**Figure S1**).

As a result of the data presented, we propose that the mechanism depicted in **Figure 5** plays a central role in magnetic field activation of TRPV1, leading to increased intracellular calcium driving downstream production of SEAP. The αGFPnb-V1/GFP-FtD magnetogenetics construct yields two related mechanisms for TRPV1 activation; one for shorter-term RF-AMF stimulation (low calcium, gray), and one for longer-term RF-AMF stimulation or RF-AMF/Cap potentiation (high calcium, green). The latter pathway involves a feedback loop leading to increased cytoplasmic Ca^2+^ levels. This pathway can be activated via long-term RF-AMF stimulation, in keeping with the greater increase in SEAP with RF-AMF at later time points, or by potentiation via capsaicin. This pathway is proposed to involve PKC activation as a result of increased Ca^2+^ levels^44^ (Lipp & Reither, 2011), which activates the p47*^phox^* subunit of NOX1/2^39^ (Cosentino-Gomes et al., 2012), leading to activation of the enzyme at the cell membrane and resulting in increased ROS levels. Inhibition of TRPV1 capsaicin potentiation by the PKC inhibitor Gö 6983 (**Figure 1E**) and inhibition of the NOX gp91*^phox^* cytochrome by GSK 2795039 (**Figure 1F**) support this hypothesis. The increased ROS can further activate TRPV1 and enhance Ca^2+^ release from the ER through IP_3_ receptors^38^ (Sakurada et al., 2019), which can be blocked by addition of Xestospongin C, an inhibitor of ER ATPase (**Figure 1C**). The increased Ca^2+^ levels ultimately increase SEAP production. Critically, in both cases, initial activation of TRPV1 by RF-AMF stimulation is driven by local ROS generation from ferritin.

**Figure 5:**
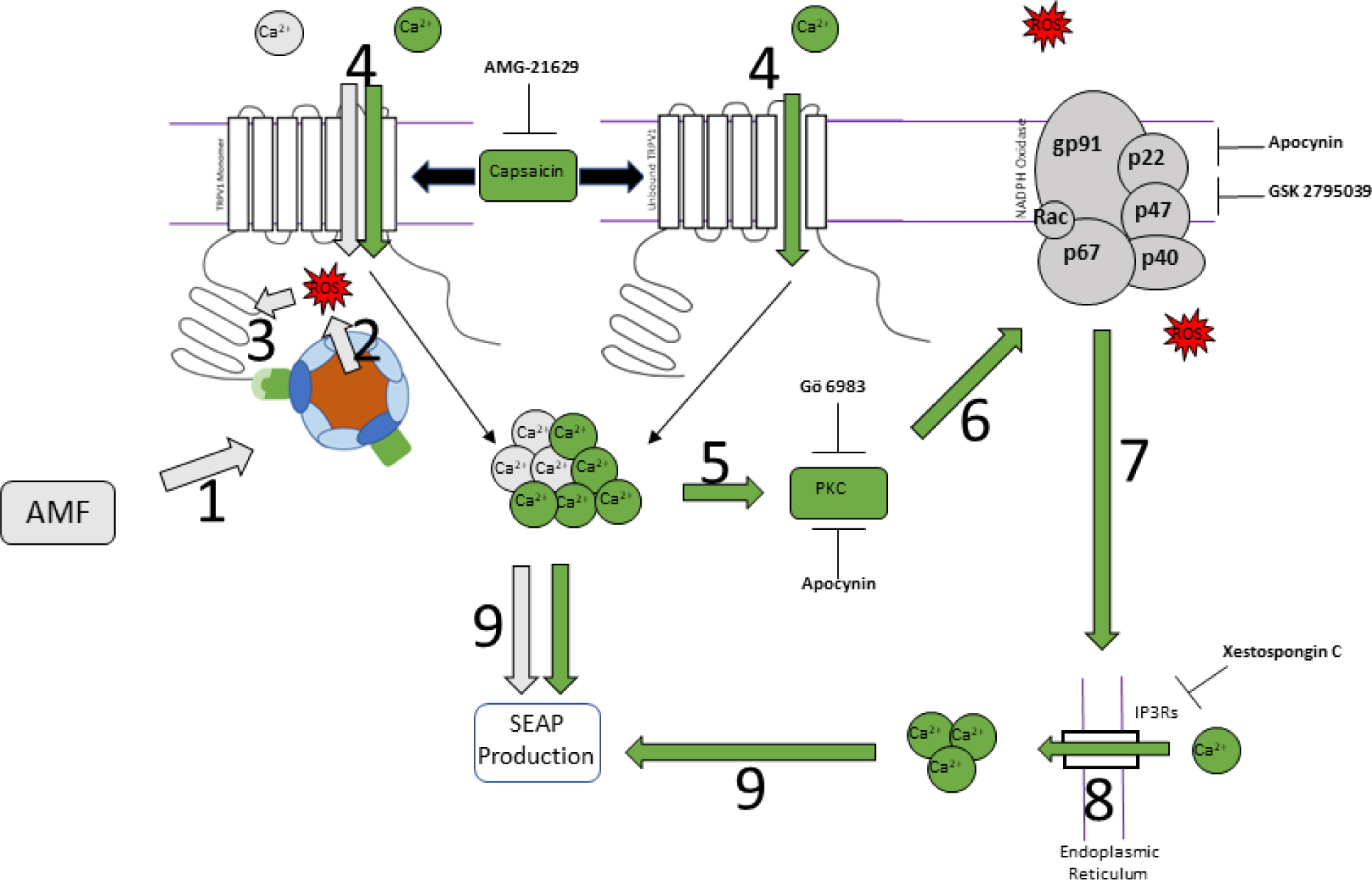
Proposed mechanism of RF-AMF driven magnetogenetics. The interrelationship of RF-AMF stimulation alone (gray) and upon potentiation with capsaicin (green). (1-2) AMF stimulation targets ferritin nanoparticles leading to local ROS generation. (3) ROS in proximity to TRPV1 reacts with crucial cysteine residues, increasing channel sensitivity. (4) Calcium gating through TRPV1 pore. (5) Heightened calcium levels lead to kinase activity, such as protein kinase C (PKC). (6) PKC activity facilitates NADPH oxidase assembly and activation in the cell membrane. (7-8) Increased NADPH oxidase activity produces additional ROS and triggers calcium release from endoplasmic reticulum through IP_3_ receptors (IP3Rs). (9) Increased cytoplasmic calcium levels lead to calcium-dependent SEAP transcription. Various inhibitors studied are shown in black.

In conclusion, the mechanism of activation of the αGFPnb-V1/GFP-FtD magnetogenetics platform is strongly associated with the generation of ROS by ferritin in the presence of an RF-AMF. Based on the current study, perhaps the most important structural requirement for RF-AMF activation of the TRPV1-ferritin system is the tethering of ferritin onto TRPV1 to ensure a sufficient ROS concentration near critical cysteine residues in the vicinity of the N- and C-termini of the ion channel. The presence of ROS localized to these cysteine residues results in their oxidation, which subsequently gates the channel. It cannot be ruled out, however, whether ROS alone is the sole driving force of RF-AMF driven TRPV1 activation or if thermal and mechanical effects still play a role. Nonetheless, it is clear that ROS plays a critical role in platform activation. Better understanding of the mechanism of magnetic field induced ion channel activation should allow for further advancements in its implementation *in vitro* and *in vivo*.

### Limitations of study

Due to the physical limitations of the RF-AMF generating unit, sample sizes are small, resulting in less-than-ideal statistical analysis and large error in some cases. Furthermore, the hypothesis regarding ROS oxidation of crucial cysteine residues within TRPV1 was only tested indirectly via engineering of the channel. For more rigorous testing, direct oxidation of the channel should be measured.

## Method Details

### Cell Lines

All cell culture experiments were performed with HEK-293T cells (ATCC, CRL-3216) grown in 10% FBS / 90% Dulbecco’s Modified Eagle Medium (DMEM; Gibco) in a 37°C / 5% CO_2_ incubator with 100% relative humidity. Maintenance flasks were split every 48-72 h. A previously generated HEK-293T cell line stably expressing αGFPnb-V1/GFP-FtD was used as well. (Brier et al., 2020).

### Gene constructs

αGFPnb-V1/GFP-FtD and calcium-dependent SEAP plasmids were the same as used previously (Brier et al., 2020). ROS-insensitive mutants were created by obtaining gblocks of TRPV1 gene containing C257S, C741S, or C257S/C741S mutations with terminal AvrII and BamHI restriction enzyme sites (Integrated DNA Technologies). Gblocks were then digested and ligated into existing plasmid using ThermoFisher FastDigest protocol. Modified genetic constructs were created by independently removing αGFPnb or GFP-FtD from the original construct. All modified constructs were confirmed via sequencing (GENEWIZ).

### RF-AMF stimulation system

RF-AMF stimulation experiments were performed using the previously described custom RF system (Brier et al., 2020). Briefly, a W-5/500 power station with HS-8 heat station was used to created 501 kHz, 27.1 mT AMFs (UltraFlex Power Technologies). Samples were placed in a custom holder capable of holding four samples at equivalent magnetic field environments within the coil. Heat generation from the coil was regulated by cold water passed through the interior of the coil. A non-CO_2_ incubator was set to the coil’s equilibrium temperature to create the corresponding AMF− temperature control condition.

Cell cultures under RF-AMF stimulation were performed as per our previous work. Briefly, HEK-293T cells and HEK-293T cells stably expressing the αGFPnb-V1/GFP-FtD were cultured as described above (same for all steps unless otherwise noted). Cells were subcultured onto 12-mm-diameter No. 2 cover glass slips coated with fibronectin (Sigma-Aldrich; 10 μg·mL^−1^ in DPBS) in 24-well plates. Cells were plated onto the cover glass slips at low density (∼40,000 cells per slip) in 500 μL of DMEM supplemented with 10% FBS. The following day, cells were transfected with the calcium dependent-SEAP construct (248 ng·well^−1^) using a standard Lipofectamine 2000 (2.5 ng:1 ng DNA; Invitrogen) protocol and supplemented with 2 mg·mL^−1^ *holo-*transferrin (Thermo 11107018). For transiently transfected experiments, HEK-293T cells were transfected with a 1:2 molar ratio of magnetogenetics construct (252 ng·well^−1^) to the CaR-SEAP construct (248 ng·well^−1^). The medium was changed the following day to 500 μL OMEM supplemented with 1% FBS and 500 μM ferric citrate, and samples were moved to a 32°C/5% CO_2_ incubator. For RF-AMF treatment, cells were transferred to 400 μL of Ca^2+^-enriched (using CaCl_2_; Sigma-Aldrich) OMEM (final Ca^2+^ concentration of 2.5 mM). To normalize data by cell number, a standard 3-(4,5-dimethyl-2-thiazolyl)-2,5-diphenyl-2H-tetrazolium bromide (MTT; Sigma-Aldrich) was employed.

RF-AMF potentiation experiments used 1.0 μM capsaicin (Tocris) in Ca-OMEM. For inhibition experiments, media was supplemented with 1.0 μM Xestospongin C (Tocris), 1.0 μM Ryanodine (Tocris), 5.0 μM Gö 6983 (Tocris), or 100 μM GSK 2795039 (Tocris). Identical samples were prepared in a 32°C incubator for the no magnetic field controls.

### ROS Measurements

Dichlorodihydrofluorescein diacetate (DCFDA) dye was used for intracellular ROS determination following Abcam’s protocol (ab113851). For extracellular measurements, 25 μM Amplex Red (ThermoFisher) and 1.0 mU/mL horseradish peroxidase (ThermoFisher) were supplemented into the cell culture media. Fluorescence was measured on a SpectraMax M5 plate reader with excitation/cutoff/emission wavelengths of 530/570/590 nm. Hydrogen peroxide (ThermoFisher) and *tert*-butyl hydrogen peroxide (ThermoFisher) were used as standards for the extra- and intracellular assays, respectively.

### SEAP Measurements

SEAP production was quantified as per our previous study (Brier et al., 2020). Briefly, SEAP production was assayed through its conversion kinetics of *para*-nitrophenyl phosphate (Sigma-Aldrich; 1 mg·mL^−1^) to *para*-nitrophenol in 1 M diethanolamine buffer (SeraCare; pH 9.8) at 405 nm, compared to an alkaline phosphatase (Roche) standard.

### Statistics

Significance was calculated using Student’s T-Test. P ≤ 0.0001 (****), 0.001 (***), 0.01 (**), 0.05 (*). Samples size of three except where noted.

## Acknowledgements

This work was supported by the National Science Foundation (CBET-1930163).

## Ethics statement

All methods were carried out in accordance with relevant guidelines and regulations. No animals were directly involved in the study.

## Data availability

The data sets generated during and that support the findings of this study are available from the corresponding author upon reasonable request.

## Author Contributions

J.W.M., M.I.B., S.A.S., and J.S.D. conceived and designed the experiments. J.W.M. and E.O. performed the experiments. J.W.M., M.I.B., S.A.S., and J.S.D. analyzed the data. J.W.M. and J.S.D. led the manuscript preparation with contributions from all authors.

## Supplemental Titles and Legends

**Figure S1: Pulsed AMF Stimulation.** HEK 293T cells stably expressing αGFPnb-V1/GFP-FtD and transfected with Ca^2+^-dependent SEAP were subjected to 50% on/off pulsed AMF over the course of 4 h (i.e., 2 h of total AMF stimulation). (A) SEAP and (B) ROS production was measured. For both, a positive correlation was seen with pulse duration. Despite total stimulation duration being the same, the increase in both ROS and SEAP with pulse duration indicates that an excess of ROS is being produced with increased AMF stimulation, allowing for continued stimulation of the platform during the ‘off’ phases. N = 3. Mean ± s.d.

**Figure S2: Inhibitor effect on ROS production.** HEK 293T cells stably expressing aGFPnb-V1/GFP-FtD magnetogenetics platform were exposed to ± AMF for 2 h with indicated inhibitor. None = Basal. Rya = Ryanodine, 1.0 μM. XC = Xestospongin C, 1.0 μM. GSK = GSK2795039 100 μM. Go = Go 6983, 5.0 μM. P < 0.05 for a, b, and c. P < 0.0001 for all + AMF over respective – AMF. N = 3 Mean ± s.d.

**Figure S3: AMF-Induced ROS.** HEK 293T cells were transfected with αGFPnb-V1/GFP-FtD magnetogenetics platform (WT) or indicated modified platforms and subjected to −/+ AMF for 2 h. All platforms showed significant increases in ROS when exposed to AMF compared to baseline conditions. The nbV1 platform experienced the smallest increase due to the removal of exogenous ferritin from the platform (endogenous ferritin unaffected). N = 3. Mean ± s.d.

**Figure S4: Capsaicin Dose Response.** HEK 293T cells were transfected with αGFPnb-V1/GFP-FtD magnetogenetics platform (WT) or indicated modified platforms and were subjected to a capsaicin gradient (0.0, 0.5, 1.0, or 5.0 μM) for 3 h at 37°C. All modified platforms showed significant response to capsaicin stimulation (p < 0.05), confirming TRPV1’s functionality in all platforms. N = 6. Mean ± s.d.

## References

1. Arnaud, C. H. (2019). Making biologics on demand. Chemical and Engineering News, 96(45), 36–40.

2. Boyden, E. S., Zhang, F., Bamberg, E., Nagel, G., & Deisseroth, K. (2005). Millisecond-timescale, genetically targeted optical control of neural activity. Nature Neuroscience, 8(9), 1263–1268. doi: 10.1038/nn1525.

3. Gossen, M., Freundlieb, S., Bender, G., Müller, G., Hillen, W., & Bujard, H. (1995). Transcriptional activation by tetracyclines in mammalian cells. Science, 268(5218), 1766–1769. doi: 10.1126/science.7792603.

4. Magnus, C. J., Lee, P. H., Atasoy, D., Su, H. H., Looger, L. L., & Sternson, S. M. (2011). Chemical and genetic engineering of selective ion channel–ligand interactions. Science, 333, 1292–1296. doi: 10.1126/science.1206606.

5. Gomez, J. L., Bonaventura, J., Lesniak, W., Mathews, W. B., Sysa-Shah, P., Rodriguez, L. A., Ellis, R. J., Richie, C. T., Harvey, B. K., Dannals, R. F., Pomper, M. G., Bonci, A., & Michaelides, M. (2017). Chemogenetics revealed: DREADD occupancy and activation via converted clozapine. Science, 357(6350), 503–507. doi: 10.1126/science.aan2475.

6. Gorostiza, P., & Isacoff, E. Y. (2008). Optical switches for remote and noninvasive control of cell signaling. Science, 322(5900), 395–399. doi: 10.1126/science.1166022.

7. Zhang, F., Vierock, J., Yizhar, O., Fenno, L. E., Tsunoda, S., Kianianmomeni, A., Prigge, M., Berndt, A., Cushman, J., Polle, J., Magnuson, J., Hegemann, P., & Deisseroth, K. (2011). The microbial opsin family of optogenetic tools. Cell, 147(7), 1446–1457. doi: 10.1016/j.cell.2011.12.004.

8. Wang, D., Tai, P. W. L., & Gao, G. (2019). Adeno-associated virus vector as a platform for gene therapy delivery. Nature Reviews Drug Discovery, 18(5), 358–378. doi: 10.1038/s41573-019-0012-9.

9. Armbruster, B. N., Li, X., Pausch, M. H., Herlitze, S., & Roth, B. L. (2007). Evolving the lock to fit the key to create a family of G protein-coupled receptors potently activated by an inert ligand. Proceedings of the National Academy of Sciences USA, 104(12), 5163–5168. doi: 10.1073/pnas.0700293104.

10. English, J. G., & Roth, B. L. (2015). Chemogenetics-a transformational and translational platform. JAMA Neurology, 72(11), 1361–1366. 10.1001/jamaneurol.2015.1921.

11. Deisseroth, K., & Hegemann, P. (2017). The form and function of channelrhodopsin. Science, 357(6356), eaan5544. doi: 10.1126/science.aan5544.

12. Kato, H. E., Zhang, F., Yizhar, O., Ramakrishnan, C., Nishizawa, T., Hirata, K., Ito, J., Aita, Y., Tsukazaki, T., Hayashi, S., Hegemann, P., Maturana, A. D., Ishitani, R., Deisseroth, K., & Nureki, O. (2012). Crystal structure of the channelrhodopsin light-gated cation channel. Nature, 482(7385), 369–374. doi: 10.1038/nature10870.

13. Nimpf, S., & Keays, D. A. (2017). Is magnetogenetics the new optogenetics? EMBO Journal, 36(12), 1643–1646. doi: 10.15252/embj.201797177.

14. Pan, Y., Yoon, S., Sun, J., Huang, Z., Lee, C., Allen, M., Wu, Y., Chang, Y. J., Sadelain, M., Kirk Shung, K., Chien, S., & Wang, Y. (2018). Mechanogenetics for the remote and noninvasive control of cancer immunotherapy. Proceedings of the National Academy of Sciences USA, 115(5), 992–997. doi: 10.1073/pnas.1714900115.

15. Lawrence, J.M., Yin, Y., Bombelli, P., Scarampi, A., Sstorch, M., Wey, L.T., Climent-Catala, A., Pixel IGEM Team, Baldwin, G.S., O’Hare, D., Howe, C.J., Zhang, J.Z., Ouldridge, T.E., & Ledesma-Amaro, R. (2022). Synthetic biology and bioelectrochemical tools for electrogenetic system engineering. Sci. Adv. 8(18), eabm5091. doi: 10.1126/sciadv.abm5091.

16. Krawczyk, K., Xue, S., Buchmann, P., Charpin-El-Hamri, G., Saxena, P., Hussherr, M.D., Shao, J., Ye, H., Xie, M., & Fussenegger, M. (2020). Electrogenetic cellular insulin release for real-time glycemic control in type 1 diabetic mice. Science, 368, 993–1001. doi: 10.1126/science.aau7187.

17. Huang, H., Delikanli, S., Zeng, H., Ferkey, D. M., & Pralle, A. (2010). Remote control of ion channels and neurons through magnetic-field heating of nanoparticles. Nature Nanotechnology, 5(8), 602–606. doi: 10.1038/nnano.2010.125.

18. Mosabbir, A. A., & Truong, K. (2018). Genetically Encoded Circuit for Remote Regulation of Cell Migration by Magnetic Fields. ACS Synthetic Biology, 7(2), 718–726. doi: 10.1021/acssynbio.7b00415.

19. Stanley, S. A., Gagner, J. E., Damanpour, S., Yoshida, M., Dordick, J. S., & Jeffrey M, F. (2012). Radio-wave heating of iron oxide nanoparticles can regulate plasma glucose in mice. Science, 336(6081), 604–608. doi: 10.1126/science.1216753.

20. Stanley, S. A., Sauer, J., Kane, R. S., Dordick, J. S., & Friedman, J. M. (2015). Remote regulation of glucose homeostasis in mice using genetically encoded nanoparticles. Nature Medicine, 21(1), 92–98. doi: 10.1038/nm.3730.

21. Stanley, S. A., Kelly, L., Latcha, K. N., Schmidt, S. F., Yu, X., Nectow, A. R., Sauer, J., Dyke, J. P., Dordick, J. S., & Friedman, J. M. (2016). Bidirectional electromagnetic control of the hypothalamus regulates feeding and metabolism. Nature, 531(7596), 647–650. doi: 10.1038/nature17183.

22. Wheeler, M. A., Smith, C. J., Ottolini, M., Barker, B. S., Purohit, A. M., Grippo, R. M., Gaykema, R. P., Spano, A. J., Beenhakker, M. P., Kucenas, S., Patel, M. K., Deppmann, C. D., & Güler, A. D. (2016). Genetically targeted magnetic control of the nervous system. Nature Neuroscience, 19(5), 756–761. doi: 10.1038/nn.4265.

23. Caterina, M. J., Schumacher, M. A., Tominaga, M., Rosen, T. A., Levine, J. D., & Julius, D. (1997). The capsaicin receptor: a heat-activated ion channel in the pain pathway. Nature, 389, 816–824. doi: 10.1038/39807.

24. Grandl, J., Kim, S. E., Uzzell, V., Bursulaya, B., Petrus, M., Bandell, M., & Patapoutian, A. (2010). Temperature-induced opening of TRPV1 ion channel is stabilized by the pore domain. Nature Neuroscience, 13(6), 708–714. doi: 10.1038/nn.2552.

25. Gaudet, R. (2008). A primer on ankyrin repeat function in TRP channels and beyond. Molecular BioSystems, 4(5), 372–379. doi: 10.1039/b801481g.

26. Ogawa, N., Kurokawa, T., Fujiwara, K., Polat, O. K., Badr, H., Takahashi, N., & Mori, Y. (2016). Functional and structural divergence in human TRPV1 channel subunits by oxidative cysteine modification. The Journal of Biological Chemistry, 291(8), 4197–4210. doi: 10.1074/jbc.M115.700278.

27. Brier, M. I., Mundell, J. W., Yu, X., Su, L., Holmann, A., Squeri, J., Zhang, B., Stanley, S. A., Friedman, J. M., & Dordick, J. S. (2020). Uncovering the role of reactive oxygen species in magnetogenetics. Scientific Reports, 10(1), 13096. 10.1038/s41598-020-70067-1.

28. Miranti, C. K., Ginty, D. D., Huang, G., Chatila, T., & Greenberg, M. E. (1995). Calcium activates serum response factor-dependent transcription by a Ras- and Elk-1-independent mechanism that involves a Ca2+/calmodulin-dependent kinase. In Molecular and Cellular Biology, 15(7), 3672–3684. doi: 10.1128/MCB.15.7.3672.

29. Grewal, S. S., Fass, D. M., Yao, H., Ellig, C. L., Goodman, R. H., & Stork, P. J. S. (2000). Calcium and cAMP signals differentially regulate cAMP-responsive element-binding protein function via a Rap-1-extracellular signal-regulated kinase pathway. Journal of Biological Chemistry, 275(44), 34433–34441. doi: 10.1074/jbc.M004728200.

30. Pan, M.-G., Xiong, Y., & Chen, F. (2013). NFAT Gene Family in Inflammation and Cancer. Current Molecular Medicine, 13(4), 543–554. doi: 10.2174/1566524011313040007.

31. Meister, M. (2016). Physical limits to magnetogenetics. ELife, 5(1), 17210. doi: 10.7554/eLife.17210.

32. Barbic, M. (2019). Possible magneto-mechanical and magneto-thermal mechanisms of ion channel activation in magnetogenetics. ELife, 8, e45807. DOI: 10.7554/eLife.4580.

33. Hernandez-Morales, M., Shang, T., Chen, J., Han, V., & Liu, C. (2020). Lipid oxidation induced by RF waves and mediated by ferritin iron causes activation of ferritin-tagged ion channels. Cell Reports, 30(10), 3250–3260. doi: 10.1016/j.celrep.2020.02.070.

34. Oka, T., Sato, K., Hori, M., Ozaki, H., & Karaki, H. (2002). Xestospongin C, a novel blocker of IP3 receptor, attenuates the increase in cytosolic calcium level and degranulation that is induced by antigen in RBL-2H3 mast cells. British Journal of Pharmacology, 135(8), 1959–1966. doi: 10.1038/sj.bjp.0704662.

35. Pisaniello, A., Serra, C., Rossi, D., Vivarelli, E., Sorrentino, V., Molinaro, M., & Bouché, M. (2003). The block of ryanodine receptors selectively inhibits fetal myoblast differentiation. Journal of Cell Science, 116(8), 1589–1597. doi: 10.1242/jcs.00358.

36. Young, L. H., Balin, B. J., & Weis, M. T. (2005). Gö 6983: A fast acting protein kinase C inhibitor that attenuates myocardial ischemia/reperfusion injury. Cardiovascular Drug Reviews, 23(3), 255–272. doi: 10.1111/j.1527-3466.2005.tb00170.x.

37. Hirano, K., Chen, W. S., Chueng, A. L. W., Dunne, A. A., Seredenina, T., Filippova, A., Ramachandran, S., Bridges, A., Chaudry, L., Pettman, G., Allan, C., Duncan, S., Lee, K. C., Lim, J., Ma, M. T., Ong, A. B., Ye, N. Y., Nasir, S., Mulyanidewi, S., Rutter, A. R. (2015). Discovery of GSK2795039, a novel small molecule NADPH oxidase 2 inhibitor. Antioxidants and Redox Signaling, 23(5), 358–374. doi: 10.1089/ars.2014.6202

38. Sakurada, R., Odagiri, K., Hakamata, A., Kamiya, C., Wei, J., & Watanabe, H. (2019). Calcium release from endoplasmic reticulum involves calmodulin-mediated NADPH oxidase-derived reactive oxygen species production in endothelial cells. International Journal of Molecular Sciences, 20(7), 1644. doi: 10.3390/ijms20071644.

39. Cosentino-Gomes, D., Rocco-Machado, N., & Meyer-Fernandes, J. R. (2012). Cell signaling through protein kinase C oxidation and activation. International Journal of Molecular Sciences, 13(9), 10697–10721. doi: 10.3390/ijms130910697.

40. Céspedes, O., & Ueno, S. (2009). Effects of radio frequency magnetic fields on iron release from cage proteins. Bioelectromagnetics, 30(5), 336–342. doi: 10.1002/bem.20488.

41. Céspedes, O., Inomoto, O., Kai, S., Nibu, Y., Yamaguchi, T., Sakamoto, N., Akune, T., Inoue, M., Kiss, T., & Ueno, S. (2010). Radio frequency magnetic field effects on molecular dynamics and iron uptake in cage proteins. Bioelectromagnetics, 31(4), 311–317. 10.1002/bem.20564.

42. Lima, E., Torres, T. E., Rossi, L. M., Rechenberg, H. R., Berquo, T. S., Ibarra, A., Marquina, C., Ibarra, M. R., & Goya, G. F. (2013). Size dependence of the magnetic relaxation and specific power absorption in iron oxide nanoparticles. J. Nanoparticle Research, 15(1), 1654. 10.1007/s11051-013-1654-x.

43. Taylor-Clark TE (2016). Role of reactive oxygen species and TRP channels in the cough reflex. Cell Calcium. 60(3), 155–62. doi: 10.1016/j.ceca.2016.03.007.

44. Lipp P, Reither G. Protein kinase C: the “masters” of calcium and lipid (2011). Cold Spring Harbor Perspectives in Biology, 3(7), a004556. doi: 10.1101/cshperspect.a004556.

