## Supplemental Figures S1-S4 for "Alternating Magnetic Fields Drive Stimulation of Gene Expression via Generation of Reactive Oxygen Species"

and Jonathan S. Dordick^1,2,4,*^

1. Department of Chemical and Biological Engineering, Rensselaer Polytechnic Institute, Troy, NY 12180.
2. Center for Biotechnology and Interdisciplinary Studies, Rensselaer Polytechnic Institute, Troy, NY 12180.
3. Diabetes, Obesity and Metabolism Institute, Icahn School of Medicine at Mount Sinai, New York, NY 10029.
4. Departments of Biomedical Engineering and Biological Sciences, Rensselaer Polytechnic Institute, Troy, NY 12180.

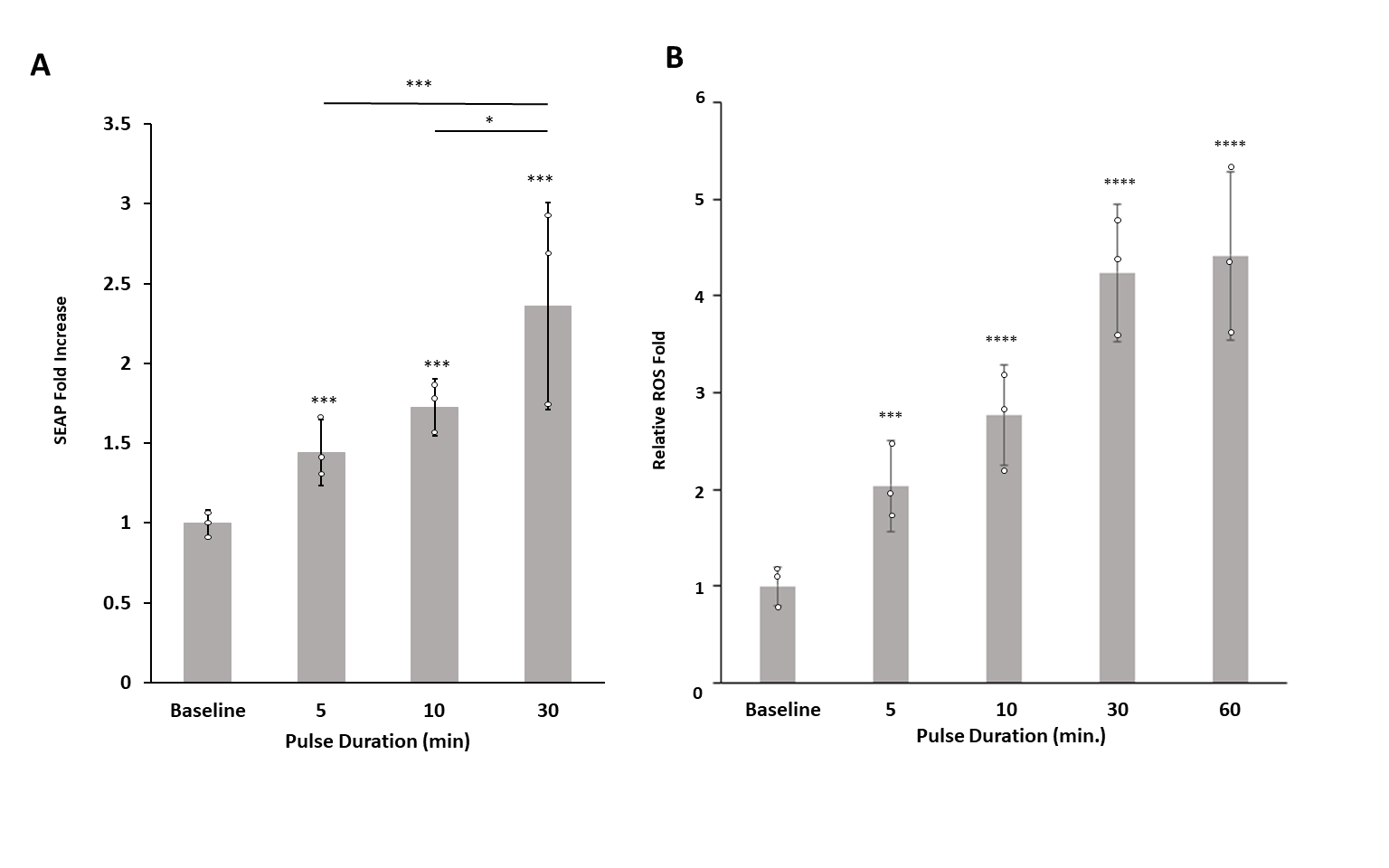

Figure S1, Mundell et al.

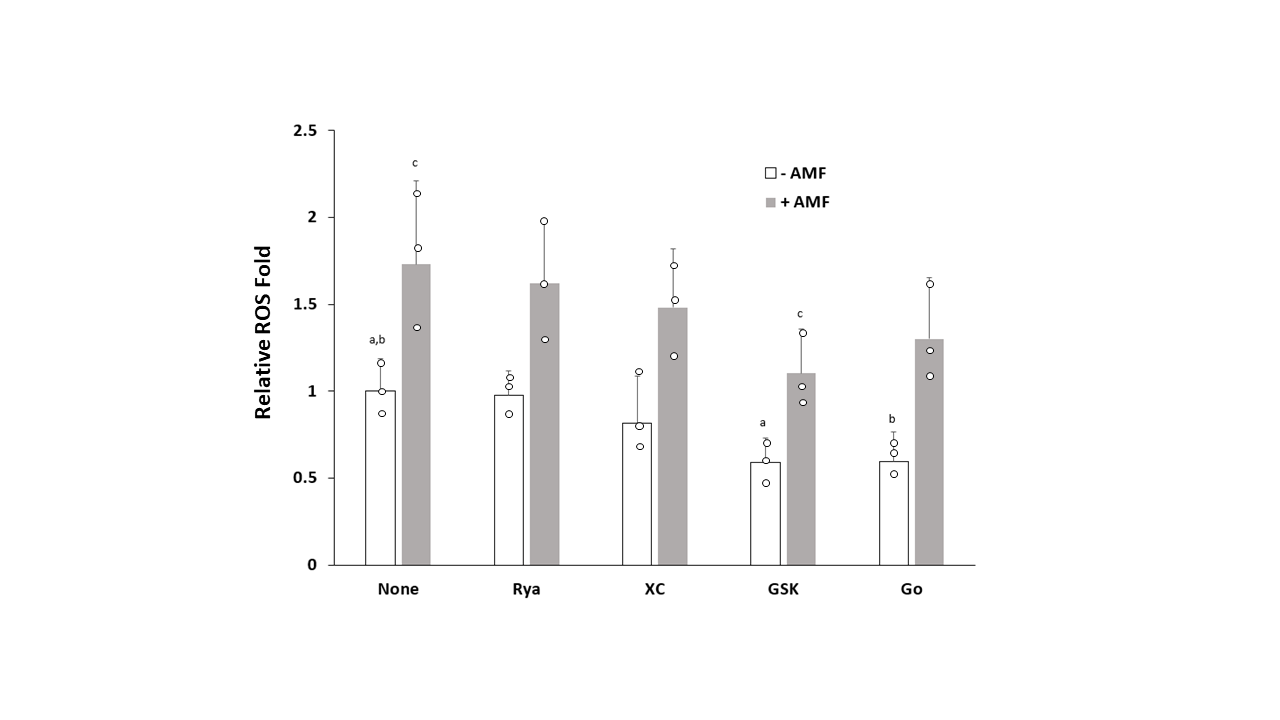

Figure S2, Mundell et al.

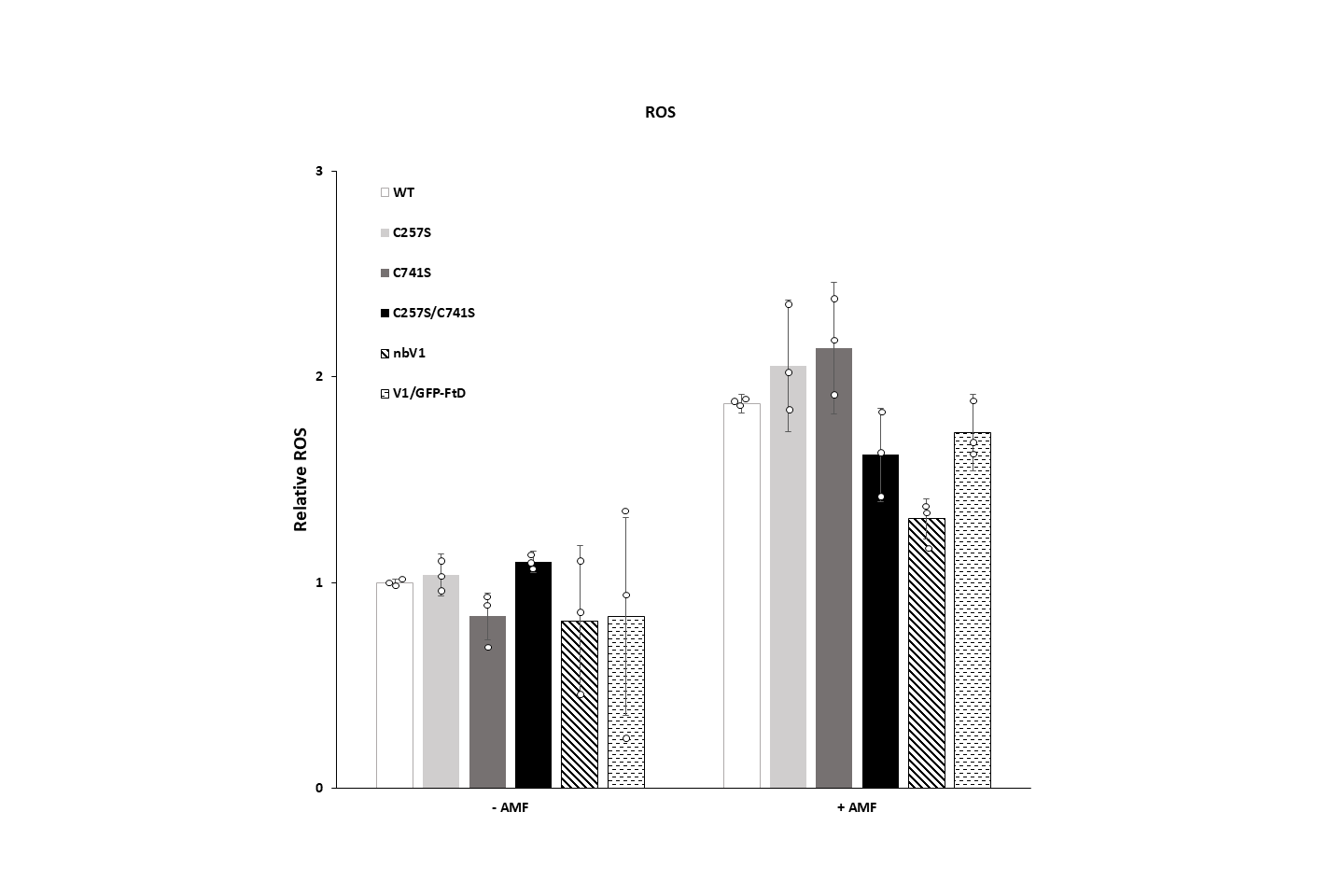

Figure S3, Mundell et al.

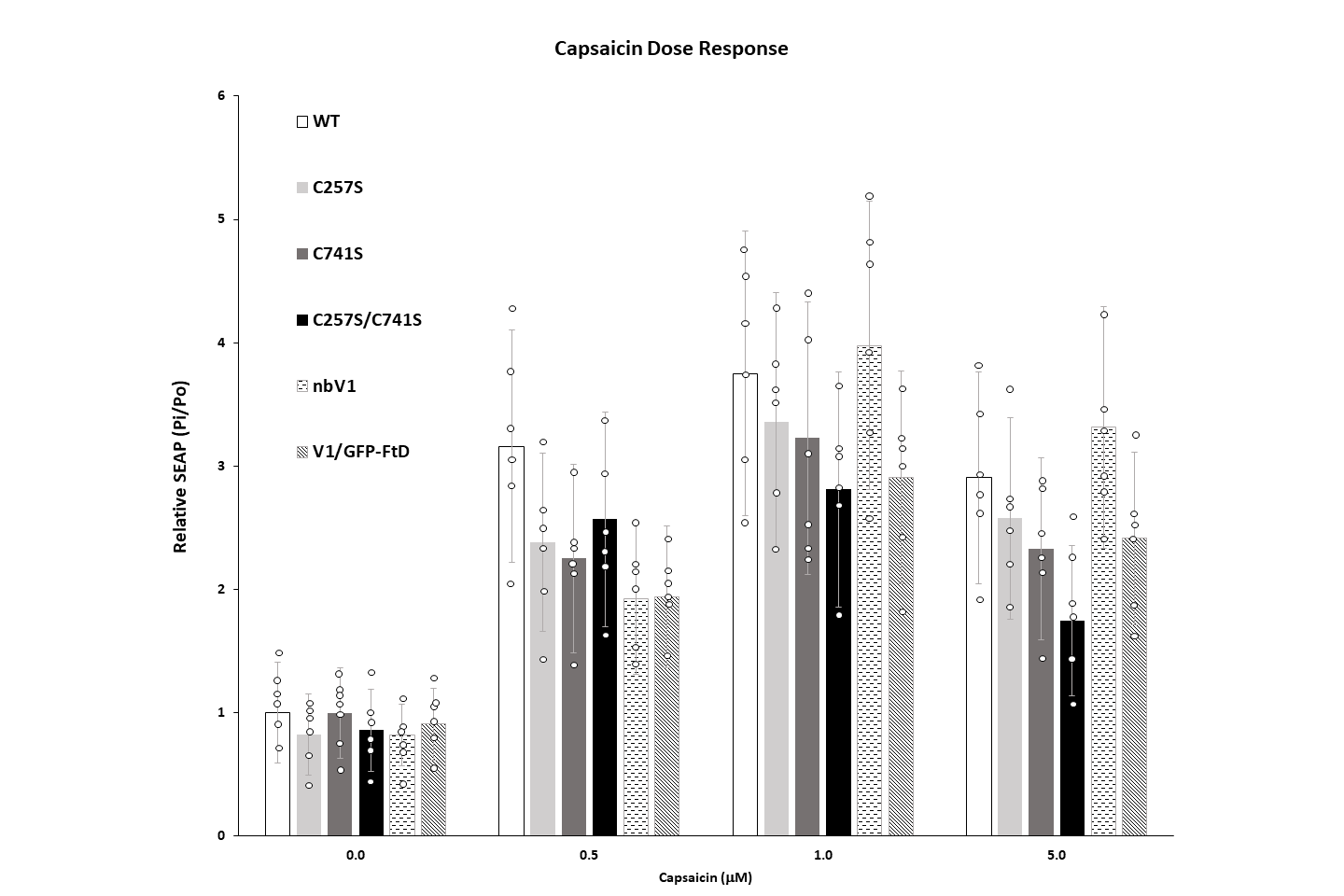

Figure S4, Mundell et al.
